# VasoTracker 2: An Open-source Platform for Quantitative Analysis of Vascular Reactivity and Function

**DOI:** 10.1101/2025.04.23.648411

**Authors:** Matthew D. Lee, Christopher Osborne, Ross Stevenson, Amy MacDonald, Grace Ebner, Danielle A. Jeffrey, Margaret A. Macdonald, Xun Zhang, Charlotte Buckley, Fabrice Dabertrand, Daniel R Machin, Jason Au, Osama F. Harraz, Nathan Tykocki, John G. McCarron, Calum Wilson

## Abstract

VasoTracker 2 is an open-source platform for studying blood vessel dynamics, featuring versatile diameter-tracking software and complementary low-cost hardware components. This system surpasses existing tools through accessible, high-resolution analysis across multiple imaging modalities, enabling comprehensive assessment of vascular dynamics in both real-time and pre-recorded experiments.

Advanced algorithms enable multi-point diameter tracking in branched vessels, automated pressure-response protocols, and reliable edge detection. The software can assess vessels imaged by brightfield microscopy, fluorescence imaging, and in ultrasound recordings, supporting diverse applications from isolated vessel studies to *in vivo* assessment.

For ex vivo applications, VasoTracker 2 includes modular open-source hardware components that can be used to create a low-cost pressure myograph system: a confocal-compatible vessel chamber and a programmable pressure controller, VasoMoto. By combining powerful analytical capabilities with an open-access approach, VasoTracker 2 provides free software and low-cost hardware alternatives to commercial systems, democratizing access to advanced vascular research tools for scientists worldwide.

## Introduction

The regulation of blood vessel diameter is a key determinant of blood flow regulation, vascular resistance and tissue perfusion. Disruptions in these control mechanisms are implicated in nearly all human diseases, either as a cause, a consequence, or as an aggravating factor. For example, reductions in vessel diameter contribute to hypertension and are linked to cardiovascular disease^1^; plaque deposits impair cerebral autoregulation in Alzheimer’s disease patients, reducing blood flow and worsening disease progression^2^; and excessive vasodilation leads to shock in sepsis patients^3^. Understanding these dysfunctional blood vessel dynamics is essential for uncovering disease mechanisms and developing effective treatments.

Quantitative analysis of vascular dynamics requires sophisticated tools that can accurately measure blood vessel diameter across diverse experimental contexts. While multiple methodologies exist for studying vascular function, ranging from isolated vessel preparations to in vivo imaging, all share a fundamental need for precise, reliable diameter measurement. Despite the critical importance of these measurements in vascular research, accessible tools for quantitative analysis across different experimental modalities have been limited. Researchers often rely on expensive commercial systems with restrictive proprietary software, or custom-developed solutions using ImageJ macros^4^ or MATLAB scripts^5,6^. This fragmented landscape raises significant barriers to entry for new researchers.

To overcome these barriers, we previously released *VasoTracker,* an open-source platform initially focused on making pressure myography - the gold standard technique for studying blood vessel regulation^7–9^ - more accessible. The initial proof-of-concept analysis software was paired with a low-cost, open-source pressure myograph that could match the performance of significantly more expensive commercial systems in assessing blood vessel diameter^10^. Whilst our initial focus was on pressure myography, the research community’s response revealed an unexpected insight: the software’s value extended far beyond our original vision. Despite its preliminary nature, VasoTracker software was adopted by vascular research laboratories worldwide and has been used to study many aspects of blood vessel physiology. For example, VasoTracker has been used to examine vasoconstriction and endothelium-dependent vasodilation^11^, the physiology of vasospam^12^, and vascular dysfunction in obesity and in cardiovascular disease^13–15^. The software has been used to study isolated blood vessel function in arteries from mice^16,17^, rats^18^, and pigs^19^. VasoTracker software has also been used to assess blood vessel function *in vivo,* in zebrafish ^20^ and in humans^21^. Moving beyond vascular studies, VasoTracker software has been used to investigate contractions in the mouse renal pelvis^22^ and bladder biomechanics^23^.

The widespread adoption and creative application of VasoTracker beyond its initial scope has driven demand for advanced features to support more diverse experimental designs. Users have requested advanced features including standalone analysis software for multiple imaging modalities, improved algorithms for complex vessel geometries, compatibility with various camera systems and automated experimental protocols. These community needs prompted the development of VasoTracker 2 – a complete reengineering of both software and hardware components. At its core, VasoTracker 2 features versatile software for blood vessel diameter tracking across multiple imaging modalities, complemented by modular hardware that, while designed in line with our original vision of pressure myography, can also be used in other systems.

Here, we introduce the new capabilities of VasoTracker 2 that address these community needs. We demonstrate how its software enables quantitative analysis across multiple imaging modalities including brightfield microscopy, fluorescence imaging, and ultrasound, and how it supports advanced applications like multi-point diameter assessment in branched vessels, and arterial strips. We also illustrate how the new modular hardware components facilitate specialized applications like automated pressure myography experiments and confocal calcium imaging in blood vessels. These advances collectively transform VasoTracker from its initial implementation into a versatile platform for investigating complex vascular dynamics in both traditional and novel experimental contexts.

## Results

VasoTracker 2 is a completely redesigned open-source, modular platform for quantitative analysis of vascular reactivity and function across multiple experimental contexts. At its core is versatile analytical software that can track blood vessel dynamics in real-time or from pre-recorded images (Figure 1), complemented by customizable hardware components for specialized applications. This integrated approach enables researchers to analyze vascular function in diverse experiments (Figure 1B) ranging from isolated vessel preparations to *in vivo* imaging.

**Figure 1.**
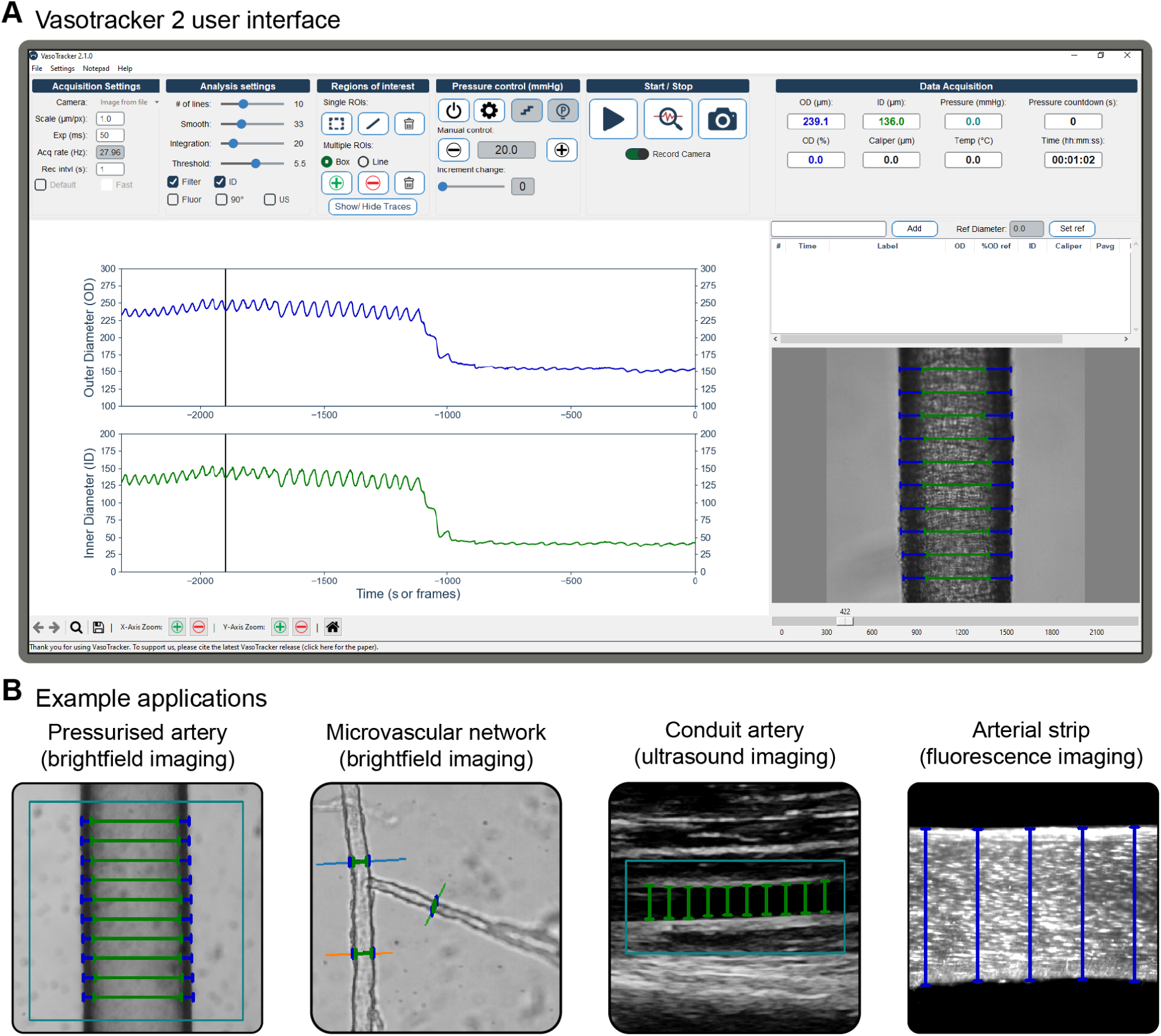
VasoTracker 2 software and tracking algorithms. A) Screenshot of the VasoTracker 2 graphical user interface. The software consists of four main panels: the settings pane, a live graph of outer and inner diameter, a table for recording experimental details, and a live or pre-recorded view of the blood vessel with diameter indicators overlaid. B) Example images showing VasoTracker diameter tracking in brightfield, fluorescence, and ultrasound modes, highlighting the versatility across different vessel preparations and imaging applications. The blue box represents a user-selected region of interest.

### VasoTracker 2 Software

The VasoTracker 2 software provides comprehensive analysis of blood vessel diameter and function across multiple imaging modalities. Built on Python 3 and µManager 2, the platform operates in both real-time and offline modes, enabling researchers to track vessel dynamics from brightfield microscopy, fluorescence, and ultrasound images. This integration with any µManager-supported camera allows VasoTracker 2 to work with existing microscope systems, making sophisticated vascular measurement techniques accessible for analyzing vascular reactivity from isolated vessel preparations to *in vivo* recordings (Figure 1B).

### Key Features of VasoTracker 2 Software

- **Multi-Modal Vessel Tracking:** Analyze vessel diameter in real-time or from pre-recorded images across brightfield, fluorescence, and ultrasound imaging modalities (Video 1).
- **Customizable Analysis Approaches:** User-defined regions or scan lines for edge detection to determine blood vessel inner diameter, outer diameter, and wall thickness with precision (Video 2).
- **Hardware Integration Capabilities:** Seamlessly integrates with VasoMoto or commercially available pressure controllers for automated experiments.
- **Universal Camera Compatibility:** Functions with any camera supported by µManager 2, eliminating hardware restrictions.

A comprehensive list of components, design files, software, and instructions for building and operating the system are available from the VasoTracker website and repository (VasoTracker and GitHub).

### VasoTracker 2 Software Applications

The original VasoTracker software^10^ established an open-source foundation for recording and analysis of bright-field images of blood vessels obtained via pressure myography, enabling live assessment of vasoconstriction, endothelium-dependent vasorelaxation, propagated vasodilation, and myogenic responses. VasoTracker 2 preserves these core functions while introducing new features that expand the range of vascular research possible with the open-source software. These include the ability to analyze pre-recorded video stacks (Figure 2A), and new algorithms for multi-point/region of interest diameter tracking in images obtained using multiple different imaging modalities. The following sections detail these new capabilities and their applications in addressing complex questions in vascular biology.

**Figure 2.**
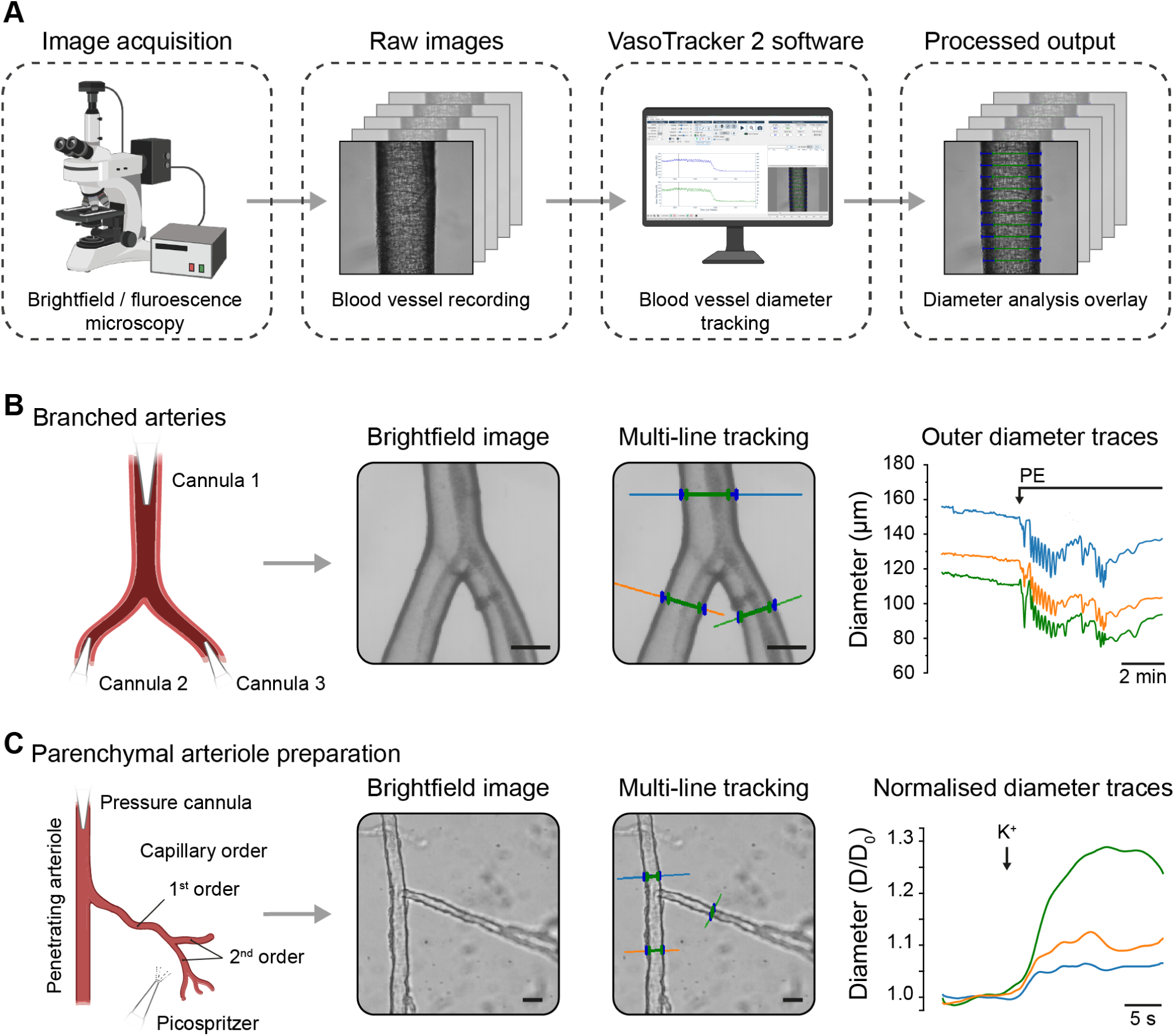
Multi-point tracking of diameter in branched blood vessels. A) Schematic of the protocol for assessing blood vessel diameter in pre-recorded images. B-C) Schematics, example images, and diameter traces illustrating the multi-line tracking feature in VasoTracker 2 in a triple-cannulated branched mesenteric artery (B) and in the capillary-parenchymal arteriole preparation (C). The data in panel B was obtained in a live experiment using fourth-fifth branch order mesenteric arteries of the rat. The third cannula was introduced to the bath and held in place using a magnetic attachment, and the artery was stimulated with phenylephrine (PE, 1 µM) to elicit vasoconstriction. Panel C shows the same algorithm applied to a pre-recording of capillary-parenchymal arteriole preparation from the mouse brain. In this experiment, 10 mM K^+^ was picospritzed onto the capillary network of the parenchymal arterioles which initiated vasodilation throughout the vascular network. Scale bars = 50 µm (B) or 20 µm (C). Components of this figure were created using bioRender.

### Multi-point Diameter Assessment in Branched Blood Vessels

The complexity of vascular networks requires simultaneous tracking of multiple vessel segments, particularly in branched structures such as cerebral parenchymal arterioles and corresponding capillary branches. The ability to track diameter changes at multiple points within these branched vessels is crucial for understanding how local and systemic factors influence the control of blood flow in the network. VasoTracker 2 introduces a multi-point tracking feature that allows concurrent assessment of diameter changes at multiple locations within a branched vessel and vascular network. This feature is shown in Figure 2, where it was applied to two distinct experimental setups.

In the first (Figure 2A), VasoTracker 2 software was used to measure the diameter of three arms of a triple-cannulated branched mesenteric artery (70 mmHg). The artery was stimulated with phenylephrine via bath perfusion, and the diameter of each branch was independently tracked during the live experiment. In the second setup (Figure 2B), VasoTracker’s multi-line tracking algorithm was applied to pre-recorded images of mouse parenchymal arterioles with attached capillary network (the CaPa preparation)^24^. In this experiment, potassium (10 mM) was picospritzed onto the capillary network. VasoTracker was used to track the diameter changes that occurred in the capillary branch, and at two locations in the parenchymal arteriole – one upstream and one downstream of the branch point.

The ability to track diameter changes in multiple branches provides insights into the coordinated vascular responses that may occur in complex networks.

### Diameter Tracking in Fluorescence Microscopy Experiments

Fluorescent labelling is commonly used in vascular studies to enhance the visualization of blood vessels. For example, fluorescent dyes permit diameter tracking whilst visualizing the internal elastic lamina separating endothelial cells and smooth muscle cells^25^ and to visualize blood vessels in vivo^26^. To demonstrate the capabilities of VasoTracker 2’s algorithms in tracking vessel diameter in fluorescently labelled blood vessels, we loaded pressurized mesenteric arteries with Calcein (1 µM) and used VasoTracker 2 to assess vascular reactivity to phenylephrine (1 µM) and acetylcholine (0.1-10 µM). Representative images and diameter traces are shown in Figure 3A, demonstrating that the algorithm accurately tracks both the inner and outer walls of the artery.

**Figure 3.**
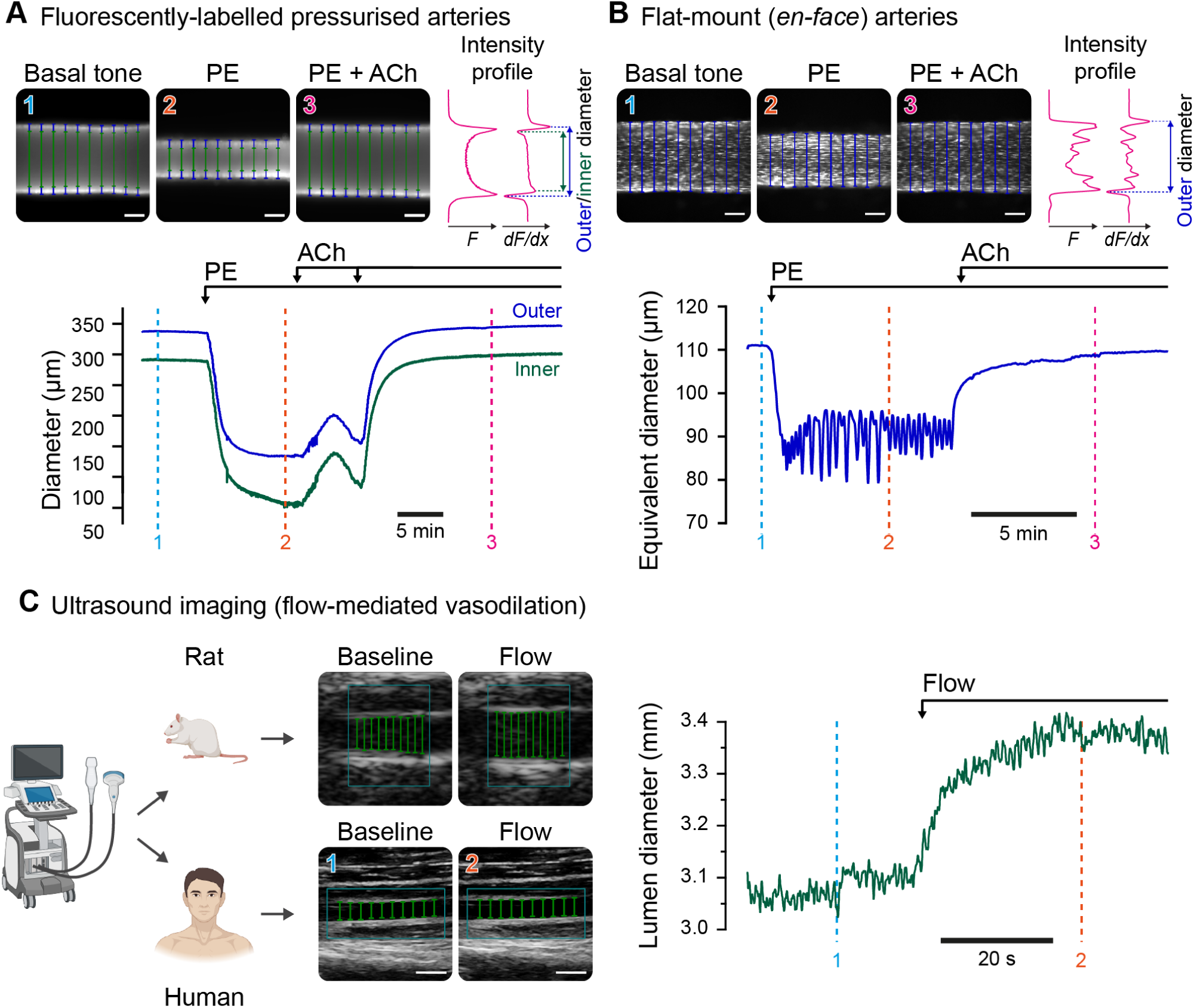
Assessing vascular reactivity using fluorescently labelled vessels and ultrasound imaging techniques. A-B) Still frame images and diameter traces of (A) fluorescently labelled pressurized arteries (Calcein, 1 µM) and (B) fluorescently labelled flat-mounted mesenteric arteries (Cal-520/AM, 1 µM). In each case, arteries were stimulated with phenylephrine (1µM) to elicit vasoconstriction, and then acetylcholine (10 nM increased to 1µM A; or 1 µM B) to induce endothelium-dependent vasodilation. The positions of artery edges are determined from intensity profiles (upper panels, F) and their derivatives (dF/dx) using the new VasoTracker 2 algorithm. When assessing flat-mounted (*en face*) arteries, inner diameter detection is prevented, and vascular reactivity is expressed as the equivalent diameter of a pressurized vessel. Scale bars = 100 µm. C) Still frame images showing flow-mediated vasodilation in rat superficial femoral and human brachial arteries. In each case, ultrasound images were collected during flow-mediated dilation experiments and stored for offline analysis using VasoTracker. Scale bars = 5 mm. D) A full trace of brachial artery diameter from a human participant from a flow-mediated dilation test. Components of this figure were created using bioRender.

Fluorescence labelling also facilitates the measurement of vascular reactivity in flat-mounted (*en face*) arterial strip preparations, a well-established method for assessing vascular reactivity^18,27–29^. Example images and a trace of vascular reactivity from a flat-mount mesenteric artery are shown in Figure 3B. In this experiment, VasoTracker 2’s algorithm was used to analyze pre-recorded images of flat-mount preparations recorded on an inverted fluorescence microscope. The images were loaded into the software and the same algorithm was employed for fluorescently labelled pressurized vessels with the exception that “inner diameter” measurement was not acquired. By incorporating these tracking algorithms into VasoTracker 2, we make fluorescence-based blood vessel tracking more accessible to the vascular research community.

### Diameter Tracking in Ultrasound Imaging

Large conduit artery diameters are most commonly measured using diagnostic ultrasound due to its widespread availability, temporal resolution for dynamic vascular responses, and non-invasive nature. Dynamic diameter tracking is particularly relevant for measures of arterial compliance^30^ and endothelial function, mostly commonly assessed via the flow-mediated dilation technique as the relative change in diameter after a shear stress stimulus^31^. Structural features of conduit arteries are also linked to future cardiovascular disease risk, for example the thickness of the intima-media complex of the common carotid artery^32^.

VasoTracker 2’s diameter tracking algorithms can be applied to ultrasound images, allowing for analysis of large conduit arteries in both animal models and clinical recordings (Figure 3C). The software’s ability to process pre-recorded images makes it suitable for retrospective analysis of ultrasound data, and allowing analysis of other datasets, providing researchers and clinicians with an open-source tool for quantitative assessment of vascular function in various cardiovascular conditions.

### Hardware Integration

Beyond tracking vessel diameter in vascular imaging datasets, VasoTracker 2 software includes features for seamless integration with hardware components. The software can communicate with various pressure control systems, including commercial controllers, traditional gravity-based setups, and the new open-source options described below. This integration enables automated experimental protocols and continuous data logging from temperature and pressure sensors, allowing researchers to precisely control experimental conditions while simultaneously tracking vessel dynamics.

### VasoTracker 2 Hardware

Building on these integration capabilities, and our continuing focus on open-source pressure myography, VasoTracker 2 includes modular hardware components for *ex vivo* applications. These components include a redesigned vessel chamber, temperature and pressure monitoring systems, and a new programmable pressure controller, VasoMoto. These modular components can be combined to create a complete pressure myograph system at a fraction of the cost of commercial alternatives (Figure 4A-B). Each component can also be used independently or integrated with existing laboratory equipment to enhance various vascular research applications.

**Figure 4.**
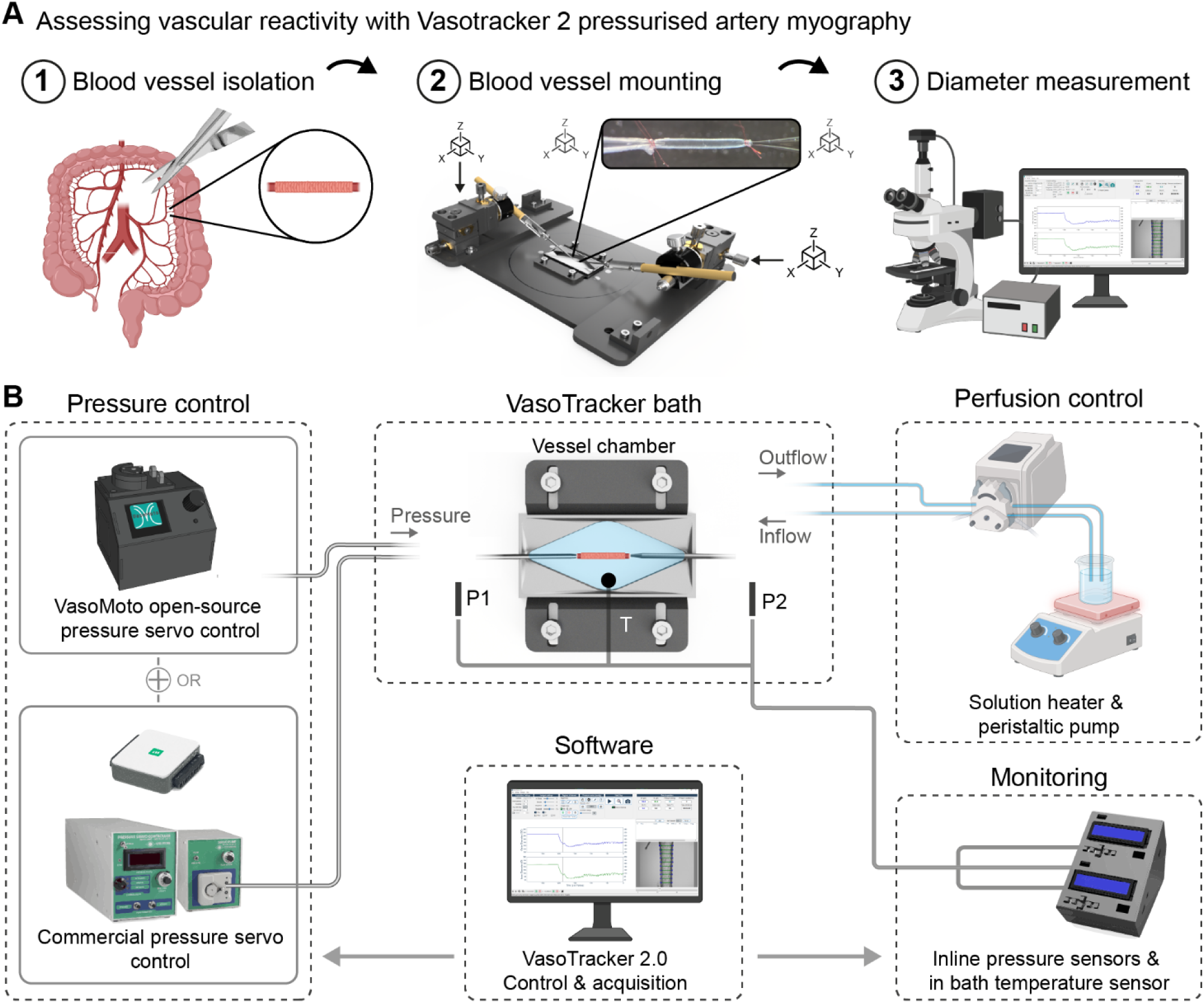
VasoTracker 2 pressure myography system. A) Schematic of the pressure myography experimental protocol. Blood vessels are freshly isolated from vascular beds of interest (1), then mounted and pressurized in the VasoTracker myograph chamber (2), before being visualized by light microscopy for diameter tracking (3). B) Schematic of the VasoTracker system. Vessels can be pressurized using the open-source, programmable pressure servo controller, VasoMoto, or commercial pressure servo controllers. Bath perfusion, temperature control, and drug delivery are managed through a peristaltic pump and a hot plate or inline heater. Inflow and outflow pressure (P1/P2), as well as temperature (T), are monitored via the Arduino-based VasoTracker 2 pressure/temperature monitor. Image acquisition, diameter measurement, and data logging are performed using the VasoTracker software. Components of this figure were created using bioRender

### Key features of VasoTracker 2 hardware

- **VasoMoto Pressure Control:** Supports manual and programmable blood vessel pressure regulation
- **Confocal-Compatible Imaging Chamber:** The chamber fits both upright and inverted microscopes.
- **Precise Cannula Positioning:** 3-axis control and angled holders for low-working distance applications such as optical sectioning.
- **Magnetic Perfusion Setup:** Quick, easy, and adjustable, attachments for fluid control.
- **Temperature Monitor:** With temperature control achieved via superfusion.

### Automated Pressure Response Curves (Myogenic Tone Experiments)

Measurement of pressure-response curves is the most common method used to investigate the intrinsic ability of blood vessels to constrict in response to an increase in pressure – myogenic tone^33^. Typically, investigators assess the ability of resistance arteries or arterioles to constrict as intraluminal pressure is increased in a stepwise fashion. The experiment is performed under both active and passive conditions to determine the degree of myogenic response at each pressure. The active conditions include calcium in the bath solution, allowing smooth muscle cells to contract, while the passive condition removes calcium to eliminate the ability of smooth muscle cells to constrict.

These experiments are labor-intensive, requiring manual adjustment of pressure and manual recording of pressure changes. To address this, we developed VasoMoto, an open-source pressure servo system and peristaltic pump specifically designed for pressure-response curve experiments. When paired with VasoTracker 2 software, VasoMoto allows complete automation of these experiments (Figure 5A, Video 3). The system can adjust intraluminal pressure stepwise at pre-determined intervals, with time and pressure changes automatically recorded in the VasoTracker graphical user interface (GUI). Beyond its cost-effectiveness compared to commercial alternatives, VasoMoto offers scalability through its modular design, allowing researchers to adapt the system for different experimental configurations (for example to deliver pulsatile pressure). Importantly, VasoTracker 2 is also compatible with commercial pressure servo systems, offering researchers the flexibility to integrate the software with their existing hardware setups.

**Figure 5.**
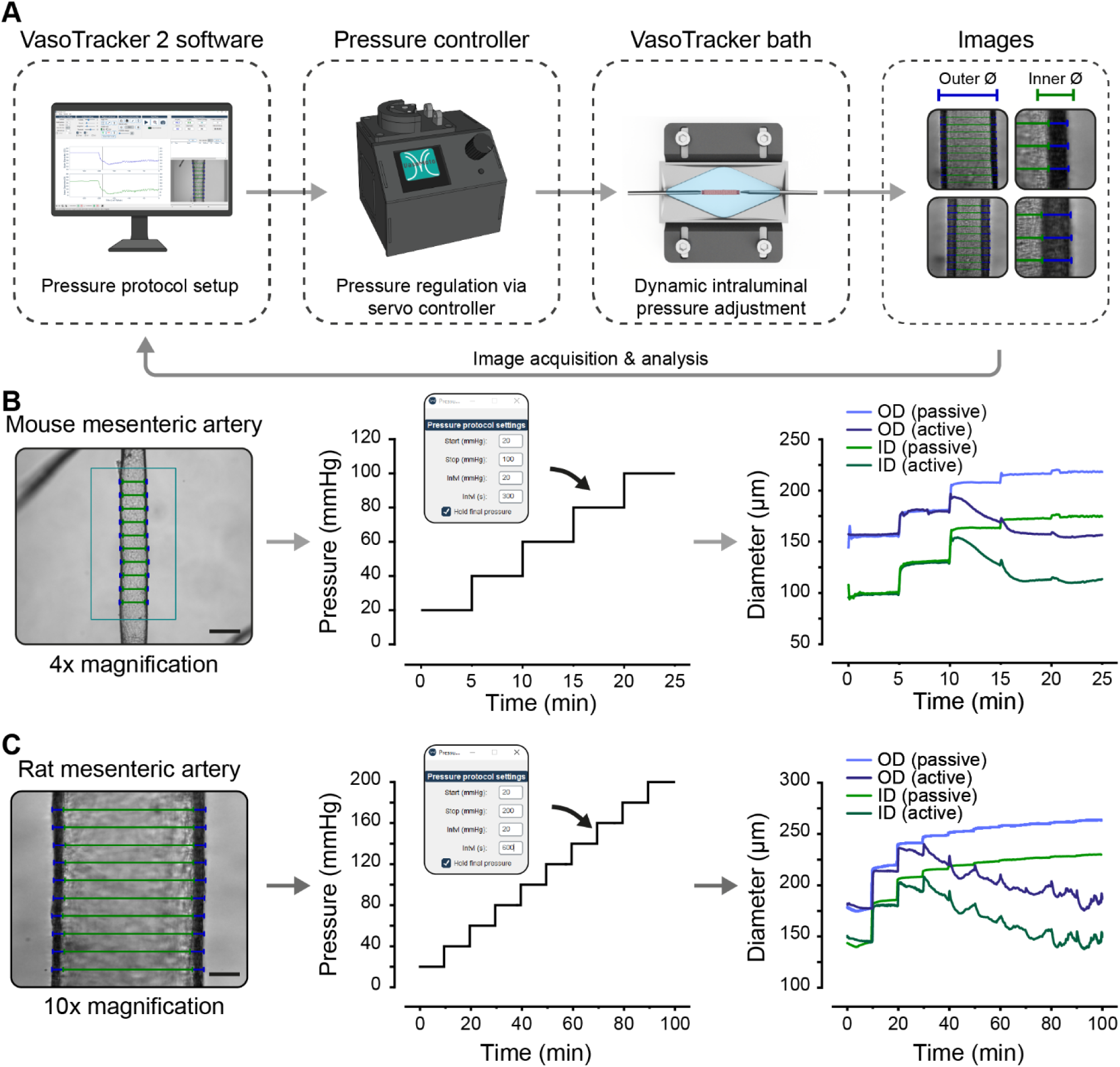
Automated myogenic tone experiments. A) Schematic of the VasoTracker 2 set up for automated control of intraluminal pressure using the open-source VasoMoto or commercial Living Systems PS-200 pressure servo systems. B-C) Example images with diameter indicators overlaid (outer diameter; blue, inner diameter; green), automatic pressure step protocols, with VasoTracker 2 pressure settings interface, and diameter traces from pressure-curve experiments performed in mouse (B) or rat (C) mesenteric arteries. Scale bars = 200 µm (B) or 50 µm (C).

Figure 5B and 5C show example data that was collected using VasoTracker automation. Myogenic responses in mouse (Figure 5B) and rat (Figure 5C) mesenteric arteries were measured using VasoTracker myographs, with pressure protocols that stepped from 20 mmHg to 100 mmHg (mouse) or to 200 mmHg (rat). Arteries from each species exhibited, and maintained, myogenic constriction at pressures above 60 mmHg. Such automation reduces the need for manual intervention, allows for more efficient data collection and increases robustness in the experiment, as the interval between each pressure increase is consistently maintained.

### Confocal Fluorescence Imaging in Pressurized Arteries

Confocal imaging is a powerful technique for studying the structure and function of blood vessels under near physiological conditions. High-resolution optically sectioned imaging can reveal detailed structural features of the vessel wall, such as the alignment of vascular cells and the expression pattern of key proteins, which are critical for understanding vascular function^34^. Optically sectioned imaging can also be used to study intracellular calcium dynamics that drive smooth muscle contraction^35,36^ and calcium signaling and nitric oxide production in endothelial cells that promote vasodilation^25,37^.

VasoTracker 2 enables these advanced imaging capabilities by being fully compatible with both upright and inverted confocal microscopes (Figure 6A-C). When used on an upright confocal microscope, the angled cannulas provide the required clearance necessary to accommodate high numerical aperture water-dipping objectives. If used on an inverted confocal microscope, the manipulators allow the vessel to be positioned close to the cover slip on the imaging chamber base so that low working distance oil immersion objectives may be used. These features allow researchers to maintain the artery under pressurized conditions while simultaneously capturing high-resolution optical sections or confocal recordings of calcium activity within smooth muscle cells (Figure 6D) or endothelial cells (Figure 6E) within the vessel wall. An additional feature of the VasoTracker 2 bath is that the modular chamber is designed so that it can replace the stage ring of commercial microscopes, making it suitable for all live cell imaging studies (Figure 6F-G).

**Figure 6.**
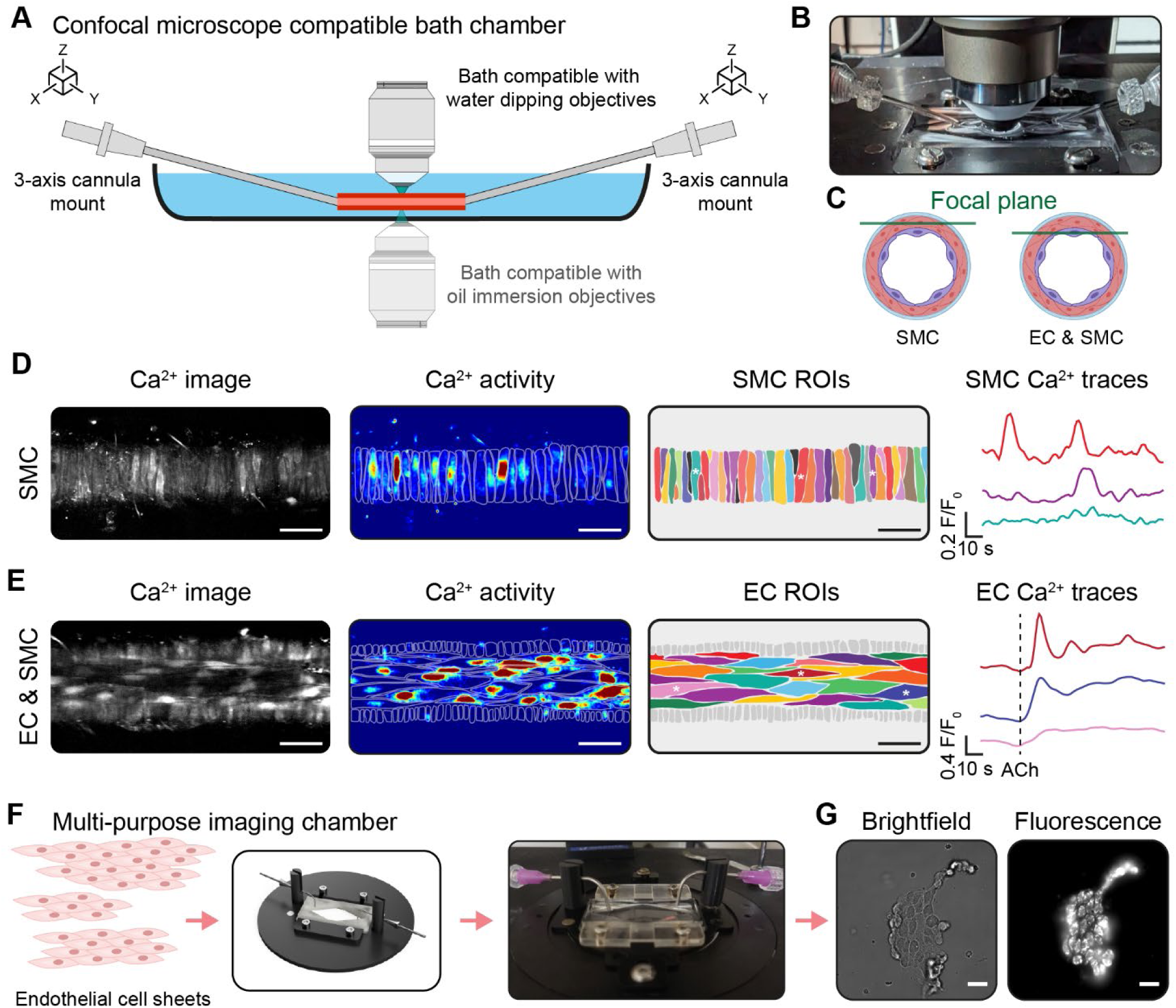
Fluorescence calcium imaging in pressurized arteries using confocal imaging. A) The 3-axis cannula mounts on VasoTracker chambers facilitate clearance for water dipping objectives on upright microscopes. Additionally, 3-axis cannula positioning allows vessels to be located close to the coverslip bottom of the chamber for compatibility with oil-immersion lenses on inverted confocal microscopes. B) Image showing the experimental setup on an upright confocal microscope. C) Diagram illustrating how optical sectioning facilitates the imaging of the smooth muscle cell (SMC) or endothelial cell (EC) layer in pressurized arteries. D-E) Representative calcium images and corresponding basal calcium signals from smooth muscle cells (D) and acetylcholine-evoked (ACh, 1 µM) activity in endothelial cells (E) in rat mesenteric arteries pressurized to 60 mmHg. F-G) Schematic (F) and example fluorescence images (G) obtained using the VasoTracker chamber for live cell imaging of acutely isolated endothelial cell sheets. The chamber is designed to replace the microscope stage ring in most commercial microscopes. Cells were isolated from rat mesenteric arteries and loaded with the mitochondrial membrane potential sensitive indicator, Mitotracker red (100 nM). Scale bars = 50 µm (D&E) or 20 µm (G).

## Discussion

Tracking blood vessel diameter is fundamental to understanding vascular function in health and disease. Changes in vessel diameter regulate blood flow, vascular resistance, and tissue perfusion, while disruptions in these mechanisms contribute to conditions such as hypertension, diabetes, neurodegenerative diseases, and sepsis. While researchers employ various methodologies to study vascular dynamics, from isolated vessel preparations to *in vivo* imaging, all require accurate diameter measurement to generate meaningful insights.

VasoTracker 2 addresses this core need through versatile blood vessel diameter tracking software that works with data from multiple imaging modalities. By supporting both real-time tracking and offline analysis of brightfield, fluorescence, and ultrasound images, the platform facilitates diverse applications from pressure myography to *in vivo* imaging studies. We have demonstrated how VasoTracker 2 can be used to assess vascular tone in fluorescent-labelled arterial strips^18^ and in conduit arteries visualised using ultrasound imaging. But the tracking algorithms could also be applied to other functional imaging datasets beyond traditional approaches, such as *in vivo* two-photon fluorescence imaging^26^, while maintaining a standardized analysis across the field.

The open-source nature of VasoTracker 2 software eliminates the need for expensive proprietary acquisition and analysis tools, democratizing access to sophisticated vascular measurement techniques. Since the software is built on μManager 2, it integrates with a wide range of cameras and microscope setups, allowing researchers to enhance existing systems without significant additional investment. This accessibility is particularly valuable for researchers in resource-limited settings and for educational institutions training the next generation of vascular physiologists.

For researchers conducting *ex vivo* blood vessel experiments, VasoTracker 2 offers additional benefits through its complementary hardware components. The system provides a fully open-source, low-cost alternative to commercial pressure myography setups. Researchers can construct our second generation pressure myography system at a significantly reduced cost of commercial alternatives. Beyond cost considerations, VasoTracker 2 offers flexibility, allowing users to tailor the system to their specific experimental needs. The redesigned vessel chamber facilitates advanced imaging techniques including confocal microscopy, whilst VasoMoto VasoMoto, a new open-source peristaltic pump and pressure servo controller, enables automated intraluminal pressure regulation, which may improve experimental reproducibility. Importantly, the modular design allows all VasoTracker 2 components to be used independently, providing researchers with tools for vessel imaging and analysis that can be adapted to a wide range of experimental applications beyond pressure myography.

The technical complexity of vascular research techniques represents another barrier that VasoTracker 2 software and hardware addresses through design improvements focused on usability and functionality. In our teaching laboratories, undergraduate students with brief training have successfully used the software for the analysis of vascular pharmacology experiments, while in our research laboratory, the complete pressure myograph system has been used by students for undergraduate research projects. Recent advancements in guidelines for assessing blood vessel function have provided standardized procedures that, combined with accessible tools like VasoTracker 2, may help new users apply these techniques effectively^8^.

Importantly, VasoTracker remains an open-source project, which facilitates continued development and community-driven improvements. By making both software and hardware designs freely available, the research community can contribute to its ongoing development. The adoption of VasoTracker by multiple laboratories indicates the value of accessible research tools, and VasoTracker 2 builds on this foundation with expanded functionality and flexibility.

While VasoTracker 2 addresses many challenges in vascular research, limitations remain. While the software works well for most vessel types, extremely small vessels (< 10 μm) or vessels with low contrast may present tracking challenges. Additionally, the hardware components currently require some, albeit minimal, technical assembly skills, and future developments could focus on further simplifying construction. These limitations provide opportunities for future improvements as the open-source community continues to develop the platform.

In summary, VasoTracker 2 represents a community-driven approach to vascular research that democratizes access to sophisticated analysis techniques. By providing versatile software for quantitative vascular analysis across multiple imaging modalities, complemented by modular, low-cost hardware components, the platform reduces barriers to entry for vascular research and has the potential to accelerate scientific discovery and foster reproducibility across laboratories worldwide.

## Methods

VasoTracker 2 is a modular open-source system for measuring blood vessel diameter, with versatile blood vessel tracking software, and complementary hardware that supports pressure myography applications. The software and hardware components each function independently, allowing researchers to integrate selected elements into existing setups or combine them into a comprehensive pressure myography solution.

### VasoTracker 2 Software

The VasoTracker 2 software is an advanced tool for tracking the diameter of blood vessels in pressure myography experiments and beyond (Figure 2). It offers enhanced capabilities for tracking diameter under a range of experimental procedures (brightfield and fluorescence imaging) and for recording user interventions. Implemented in Python 3, VasoTracker 2 is built on Python bindings of the C++ core of the μManager microscopy control system^38,39^. The software employs a modular Model-View-Controller (MVC) architecture linking the μManager bindings with VasoTracker’s 2 tracking algorithms and the graphical user interface (GUI), and facilitating easy extension (e.g. support for new cameras). Key software packaged include: NumPy^40^, scikit-image^41^, Matplotlib^42^, Christopher Gohlke’s Tifffile^43^, and Marcos Duarte’s Detect Peaks function^44^.

VasoTracker 2 software supports five primary functions:

- **Image acquisition and recording:** Captures and records images of pressurized blood vessels using a wide range of microscope-attached digital cameras.
- **Real-time diameter analysis:** Calculates, graphs, and records blood vessel diameter in real time.
- **Pressure control and data acquisition:** Controls pressure-servo systems via an analogue-digital converter for automatic pressure control and collects data from Arduino-based temperature and pressure sensors.
- **User input logging:** Allows users to log interventions, such as the addition of biological compounds to the vessel chamber.
- **Pre-recorded image analysis:** Facilitates diameter measurement in pre-recorded images, enabling flexible post-experiment analysis.

The GUI is split into four parts (Figure 2A): a control panel for adjusting key camera and analysis settings; an interactive live graph panel displaying auto-scrolling diameter measurements; a real-time image feed with diameter indicators; and a data entry table for logging experimental treatments. The interface allows users to configure settings, view dynamic data, and record experimental notes during live experiments. Additional settings are accessible through the File menu. Additional menus for controlling advanced settings (e.g., controlling the graph display, inputting the pressure protocol) are accessible through the program’s menu bar.

### Camera Compatibility & Settings

VasoTracker 2 software supports a wide range of digital cameras. Default support is provided for Basler and Thorlabs cameras, which can be readily configured using pre-defined settings. Additionally, VasoTracker 2 provides native support for μManager configuration files, allowing the integration of a wide variety of additional cameras. Essential exposure control is handled by VasoTracker, but other camera settings (e.g. bit depth, binning, gain) may be configured in μManager and exported to VasoTracker via the configuration file. By default, VasoTracker limits data acquisition to 5 Hz, but this may be increased to ∼15 Hz on suitably equipped computers.

### Tracking Algorithms & Regions of Interest

The original VasoTracker software measured vessel diameter in brightfield images using horizontal scanlines. VasoTracker 2 software maintains this fundamental approach - detecting vessel edges by identifying peaks in the derivative of intensity profiles – but builds upon the original algorithm by adding the ability to limit vessel tracking to 5 user-defined regions of interest or custom-drawn lines (Figure 2B). These can be oriented horizontally, vertically, or the user can track diameter in up to 5 custom lines drawn at any angle. For fluorescent samples, VasoTracker 2 introduces a new algorithm tailored to their unique intensity profiles. Such enhancements may allow users to examine multiple vessel branches simultaneously or avoid artefacts like vessel side-branches or residual adherent tissue.

### Data Export

VasoTracker software exports data in comma-separated value (CSV) format, ensuring compatibility with all common data analysis packages. Each experiment generates two CSV files: one containing diameter and sensor measurements, and the other with information that was entered into the table (e.g. user notes on drug additions, automatically controlled pressure changes etc.). Users can choose to record images, both raw and with tracking indicators overlaid; these images are saved in Tag Image File Format (TIFF). Additionally, users can add experimental notes in the built-in VasoTracker 2 notepad, which are saved as a Text File Document (TXT).

### VasoTracker 2 Hardware

To complement the software’s versatile analysis capabilities, VasoTracker 2 includes modular hardware components for researchers working with isolated blood vessels. These hardware elements can be used together to create a complete pressure myography system or integrated individually with existing laboratory equipment.

### VasoTracker 2 Pressure Myograph Chamber

The pressure myograph chamber design (Figure 1A) represents a significant departure from our original. The updated version is a modular bath consisting of a base and a low-volume (∼2 ml) vessel chamber insert. The low volume chamber is optimized for efficient use of reagents and precise environmental control via superfusion. The base can be machined from POM-C, and the chamber insert from acrylic, via any commercial CNC machining service. These materials ensure durability and compatibility with various experimental conditions. The modular base is designed to fit a Thorlabs (UK) MLS series motorized scanning stage but can be directly placed on any manual microscope stage or easily adapted for other automated stages.

The bath is equipped with dual MPH3 pipette holders (WPI Inc., USA), mounted to miniaturized 3-axis translation stages (DT12XYZ, Thorlabs, USA) via rotatable probe clamps (MCU, Siskiyou, USA). This configuration facilitates independent XYZ positioning of both cannulas, offering enhanced precision and flexibility. This setup makes it easier to align and manipulate blood vessels during mounting. It also allows blood vessels to be positioned near the bottom of the chamber, adjacent to the glass coverslip, for low-working distance or oil objective use on inverted microscopes. Additionally, the angled cannula design provides clearance for water dipping objectives on upright microscopes. Superfusion plumbing can be securely and flexibly attached using magnetic holders

### Pressure Control

Three options are provided for pressure control. The first is VasoMoto, a newly developed manual pressure servo system. VasoMoto is a low-cost, open-source, 3D printed peristaltic pump specifically designed for pressure myography. VasoMoto consists of an Arduino-based microcontroller, a custom instrumentation amplifier with a 16-bit analog-to-digital converter, and a precision stepper motor. The enclosure and pump head are 3D printed with minimal additional hardware. This unit interfaces directly with VasoTracker 2 to share continuous pressure measurement. VasoMoto is also capable of oscillating pressure to mimic pulsatile pressure changes seen *in vivo* to rates as fast as the murine heart rate (∼400 oscillations per minute). Since Arduino-based microcontrollers can directly interface with a computer without the need for additional hardware, VasoTracker 2 can acquire pressure data without the need for an external interface or additional software.

VasoTracker 2 also supports automated pressure regulation using PS-200 Pressure Servo Controllers (Living Systems, USA), which are programmable via a National Instruments DAQ system for precise control when intraluminal flow is not required. Finally, VasoTracker remains compatible with traditional methods that employ height-adjustable reservoirs connected to the glass cannulas in the myography bath and uses gravity to generate create a pressure gradient across the vessel when required for experiments with luminal flow^10^.

### Temperature & Pressure Monitor

The VasoTracker 2 system combines temperature and pressure monitoring. The chamber’s temperature is continuously monitored by a 10k NTC Thermistor, with real-time readings displayed on an LCD and reported to the VasoTracker software. When pressure is managed using VasoMoto, pressure readings are transmitted directly to the VasoTracker software. When height adjustable reservoirs are used (for experiments with luminal flow), pressure levels may be monitored by flow-through pressure transducers, providing real-time pressure readings on an LCD screen and transmitting data to the VasoTracker software.

### Choice of Imaging System

The myograph chamber is designed for versatility and is compatible with most microscope setups. Data presented here were acquired using various microscopes, including Eclipse Ts2R (Nikon, Japan) and CK40 (Olympus, USA) inverted microscopes equipped with 4x and 10x objectives and CMOS cameras (CS165MU/M, Thorlabs, USA; or ACE a2A1920-160umBAS, Basler, USA), and an FN-1 upright microscope with a 10x water-dipping objective (Nikon, Japan).

### Experimental Protocols

To assess performance, we tested the VasoTracker 2 software across a range of experimental conditions and imaging modalities. The software was evaluated in both real-time vessel tracking with the VasoTracker 2 pressure myograph and offline image analysis of datasets generated using other experimental techniques using brightfield, fluorescence, and ultrasound imaging.

### Animals

All experiments described in this study involving arterial samples were conducted using animals with appropriate ethical approval. Arteries obtained from male (8–12-wk-old) Sprague-Dawley rats, with all procedures approved by the University of Strathclyde Animal and Welfare Ethical Review Committee and conducted in accordance with UK Home Office regulations (Animals (Scientific Procedures) Act, 1986). Animals were euthanized by cervical dislocation with secondary confirmation via decapitation in accordance with Schedule 1 of the Animals (Scientific Procedures) Act 1986. Some experiments included male mice in Michigan and female mice in Vermont (indicated in respective figure legends). Mice were euthanized by intraperitoneal injection of sodium pentobarbital (100 mg/kg) followed by immediate decapitation. In these experiments, all procedures involving these animals were performed in accordance with local ethical guidelines and received the necessary institutional approvals (University of Colorado, Anschutz Medical Campus Institutional Animal Care and Use Committee, or the Institutional Animal Care and Use Committee of the University of Vermont). In addition, ultrasound experiments were performed on anesthetized rats at the University of Utah as previously described^45^. These procedures were approved by the University of Utah and Salt Lake City Veterans Affairs Medical Center Animal Care and Use. This study complies with ARRIVE guidelines.

### Humans

All human data presented in this study are collected from healthy human participants who provided informed written consent prior to data collection. The acquisition protocols were approved by the University of Waterloo ethics board (ORE 22477) and complied with the Declaration of Helsinki regarding the use of human participants, except for registration in a database.

### Pressurized Artery Myography

Following euthanasia, mesenteric arcades were removed and transferred to a physiological salt solution (PSS) of the following composition (mM): 125.0 NaCl, 5.4 KCl, 0.4 KH_2_PO_4_, 0.3 NaH_2_PO_4_, 0.4 MgSO_4_, 4.2 NaHCO_3_, 10.0 HEPES, 10.0 glucose, 2.0 sodium pyruvate, 0.5 MgCl_2_, 1.8 CaCl_2_ (adjusted to pH 7.4 with NaOH). Third and fourth order mesenteric artery segments were mounted in a VasoTracker 2 pressure myograph and visualized via light microscopy, at 60 mmHg (unless otherwise indicated) and 37 °C. For confocal calcium imaging experiments, arteries were loaded with the calcium indicator, Cal-520/AM (5 μM) and visualized using an Aurox Clarity structured illumination LED confocal imaging system. In other experiments, arteries were loaded with Calcein (1 µM) and visualized via epi-fluorescence microscopy using LED illumination (PE-4000; CoolLED, UK). One set of experiments examined vascular reactivity in flat-mounted (*en face*) arteries. In these experiments, arteries were cut open and the endothelium was preferentially loaded with Cal-520/AM and visualized using wide-field epi-fluorescence microscopy. In experiments examining cortical capillary-parenchymal arterioles preparations, experiments were performed as described previously^24^. Data from the flat-mounted arterial preparation and the parenchymal arteriole preparations were analyzed using the offline analysis feature of VasoTracker 2.

### Flow-Mediated Dilation

**Animals:** Rats were anesthetized under 3% isoflurane and 100% oxygen in a closed chamber anesthesia machine for approximately 1-3 min. Anesthesia was maintained using a nose-cone, and rats were secured in a supine position on a heated examination table to maintain body temperature that was equipped with ECG (VisualSonics, Toronto, ON, CAN). Briefly, an occlusion cuff (Harvard Apparatus, Fairfield, NJ, USA) was placed proximal to the right ankle. The ultrasound Doppler probe was placed proximal to the occlusion cuff on the superficial femoral artery. Measurement of superficial femoral artery vessel diameter was performed using a high-resolution micro-ultrasound imaging system (VisualSonics, Toronto, ON, CAN) equipped with an ultra-high frequency linear array transducer operating at 30-70 MHz (VisualSonics, Toronto, ON, CAN). The sample volume was maximized according to vessel size and was centered within the vessel on the basis of real-time ultrasound visualization. The superficial femoral artery was insonated and measurements of diameter were recorded at rest. After which, the occlusion cuff was inflated to occlude the distal tissue for 5 minutes. Artery diameter was recorded continuously throughout the occlusion period and for 3 minutes after cuff release. End-diastolic, ECG R-wave-gated images were collected from the ultrasound machine for off-line analysis of superficial femoral artery vasodilation using VasoTracker 2.

**Humans:** The brachial artery of a 21-year old female was imaged in B-mode with an 9-12 MHz linear array ultrasound probe (Vivid iq; GE Healthcare; Chicago, IL, USA) in the longitudinal plane (3 cm depth, 32 Hz frame rate). A pneumatic cuff positioned on the distal forearm was inflated to an occlusion pressure of 200 mmHg (minimum of 50 mmHg above systolic blood pressure) for 5 min, and subsequently released while images were continuously collected for an additional 3 min during reactive hyperaemia and stored for subsequent offline analysis using VasoTracker 2.

## Acknowledgements

This work was funded by the British Heart Foundation (RG/F/20/110007; PG/20/9/34859), and funding from The Strathclyde Institute of Pharmacy and Biomedical Sciences whose support is gratefully acknowledged. CO’s work on the project was provided by the University of Strathclyde’s Faculty of Science Research Software Engineering group. FD is supported by 2 research grants from the Ludeman Family Center for Women’s Health Research located at the University of Colorado Anschutz Medical Campus (2019 and 2024); 2 research grants from the University of Pennsylvania Orphan Disease Center in partnership with the cureCADASIL (2019 and 2022), a research grant from theLeducq Foundation for Cardiovascular Research (Leducq Transatlantic Network of Excellence 22CVD01 BRENDA), the National Institute of Neurological Disorders and Stroke (NINDS; RF1NS129022 and RF1NS140137), and the National Heart, Lung, and Blood Institute (NHLBI; R01HL136636). DAJ is supported by the National Heart, Lung, and Blood Institute (5T32GM007635 and F31HL170645). OFH is supported by the National Heart, Lung, and Blood Institute (NHLBI; R01HL169681), the National Institute on Aging (NIA; R21AG082193), the National Institute of General Medical Sciences (NIGMS; P20GM135007), the American Heart Association (20CDA35310097). the Bloomfield Early Career Professorship in Cardiovascular Research, the Totman Medical Research Trust, and the Cardiovascular Research Institute of Vermont. NRT is supported by the National Heart, Lung and Blood Institute (P01HL152951) and the National Institute of Diabetes and Digestive and Kidney Diseases (R01DK119615 and R01DK135696). DRM is supported by the National Center for Complementary and Integrative Health (NCCIH; R00 AT010017). We also thank the VasoTracker community for their ongoing contributions to the project.

## Data availability

Data supporting the findings of this study are available for download with the VasoTracker software, or from the corresponding authors on request. All code and design files are available via the VasoTracker website and associated repositories (https://www.vasotracker.com/).

## Author Contributions

CW, JGM & ML developed the concept. CW, MDL, and CO wrote the VasoTracker software, while CW and NT developed the Arduino code. CW and MDL designed the experimental apparatus. CW, ML, AM, JA, DJM, RS, GE, FB and DAJ performed the experiments. DAJ, NT, FD and OH were alpha testers and contributed valuable feedback. CW drafted the manuscript. All authors revised, edited, and approved the final version of the manuscript.

